# Does Functional Recovery Imply Stable Circuitry in the Spinal Animal?

**DOI:** 10.1101/2022.09.09.507338

**Authors:** Omar Refy, Hartmut Geyer

## Abstract

Spinal animals can regain locomotor function through gait training. However, the neural processes involved in this recovery are poorly understood. Here we use computer simulation to address if the reorganization of spinal circuits associated with the functional recovery leads to meaningful, stable circuitry function. Specifically, we develop a neuromuscular model of a spinalized rat whose circuitry can adapt based on two alternative Hebbian learning strategies, one designed to guide the circuitry back to its normal pre-injury state and the other designed to destabilize it and drive it into saturation. Exposing the model to simulated gait training, we find that both strategies lead to recovery of locomotor function as defined by the outcome measures reported in studies with spinal rats. If anything, the results obtained with the destabilizing learning strategy seem to agree more with animal observations, since it produces similarly excessive amplitudes in muscle activity. Our results suggest that gait training of spinalized animals does not necessarily effect a meaningful recovery of their spinal circuitry function. More experimental work should be directed to clarify this point, as it may have grave implications for the potential of gait rehabilitation in patients with motor complete injuries of the spinal cord.

## 1. Introduction

An animal is said to be spinalized when the neural connections between its brain and spinal cord are severed by trauma, surgery or otherwise. Although such an injury paralyzes the lower limbs, it has repeatedly been shown in experiments with spinalized cats and rats that gait training augmented by electrical stimulation and pharmacological intervention below the site of the lesion can recover some locomotor function of the hind limbs (Barbeau and Rossignol, 1987; Leon et al., 1999; Rossignol et al., 2001; Ichiyama et al., 2005, 2008; Courtine et al., 2009; Lavrov et al., 2008; Moraud et al., 2016). Yet it remains largely unknown how the nervous system achieves this functional recovery. Researchers agree that neural plasticity within the spinal cord is essential (Leon et al., 1999; Rossignol and Frigon, 2011), that electrical and chemical stimulations help to activate this plasticity (Courtine et al., 2009; Kiehn, 2016), and that it seems to reorganize nonfunctional spinal circuits into functional ones under the influence of afferent information (Dietz, 2003; Courtine et al., 2009). However, it remains unclear what the neural mechanisms are that cause this reorganization. Nor is it clear whether these mechanisms can truly effect a meaningful recovery of spinal circuit function in the absence of supervision by the brain.

We seek the help of computational modeling to address the second point. Computer models have a long tradition in the study of spinal locomotion, mainly to infer the circuitry involved in its control including muscle synergies, pattern generators, and muscles reflexes (Taga et al., 1991; Orjan Ekeberg and Pearson, 2005; Prochazka and Yakovenko, 2007; Geyer and Herr, 2010; Aoi et al., 2013; Dzeladini et al., 2014; Song and Geyer, 2015; Van Der Noot et al., 2015; Fujiki et al., 2018; Aoi et al., 2019). More closely related to functional recovery, some recent models have combined the neuromuscular locomotor apparatus with finite element representations of the lumbar spine to better understand how electrical stimulation affects the reflex circuitry in spinalized animals (Capogrosso et al., 2013; Moraud et al., 2016). In contrast, we focus on the functional reorganization itself, and present a neuromuscular model of a spinalized rat whose neural circuitry adapts during gait training. To address if the functional recovery observed in animal experiments implies a meaningful recovery of spinal circuit function, we implement two alternative variants of Hebbian learning for this circuit adaptation. The first variant is designed to guide the circuitry towards its meaningful pre-injury state (stabilizing), whereas the second one simply drives the circuitry into saturation (destabilizing). We find that despite a stark contrast in the resulting circuit structure, both implementations lead to recovery of gait as reported in the literature. Our results suggest that the current experimental observations do not suffice to determine if functional recovery of spinal animals reflects a meaningful recovery of circuit function, and we discuss potential experiment modifications that could help to resolve this point.

## 2. Simulated Gait Training of Spinal Rat

To compare model predictions against outcomes of functional recovery reported in the literature, we simulate gait training of a spinal rat as performed in animal experiments (Rossignol et al., 2001; Ichiyama et al., 2008; Courtine et al., 2009). The simulation comprises a lower body musculo-skeletal model of a spinalized rat controlled by spinal circuits and exposed to gait training on a treadmill while being partially suspended in a body-weight support (BWS) system (Fig. 1).

**Figure 1.**
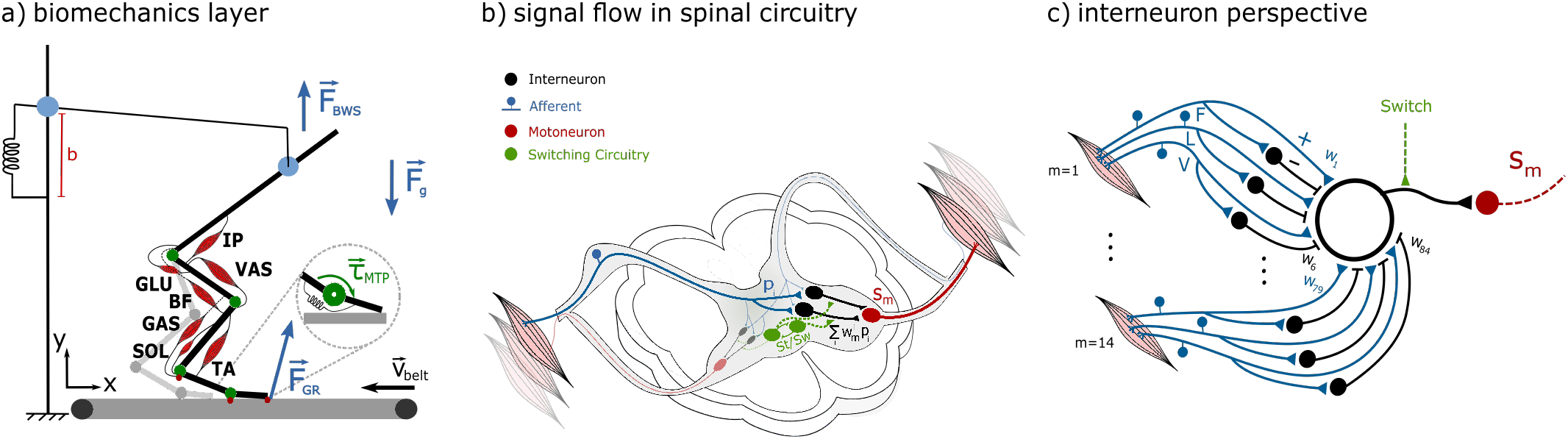
Neuromuscular model of a spinal rat that undergoes body-weight-supported treadmill training. **a)** Musculoskeletal system and training environment. The rat model comprises a trunk connected to two hind limbs in the sagittal plane. Each limb has four segments actuated by seven muscles at the hip, knee, and ankle joints. An additional MTP joint is passively actuated by a spring-damper force. Foot contact points (indicated by red dots on the bottom of the foot) create ground interaction forces *F_GR_* on the treadmill belt surface, which moves with speed *ν_belt_*. The body suspension system is modeled as a damped spring in the vertical direction that supports the trunk with *F_BWS_* and mechanical constraints at the suspension point that prevent motion in other directions. **b)** Example flow of a single signal in the model’s spinal circuitry. An afferent signal (blue) projects into interneuronal circuitry (black), which generates input to a contralateral motoneuron (red) that stimulates muscle. For each muscle, there exist two interneurons, one for stance control and another for swing control. Switching circuitry alternates between these two. **c)** Each interneuron receives afferent input from all 14 muscles of the rat model. These muscle each generate six input signals representing positive (+) and negative (-) afferent feedbacks based on muscle force (F), muscle fiber length (L) and muscle contraction velocity (V). Thus, in total an interneuron receives 84 input signals modulated by the synaptic weights *w*_1_... 84.

### 2.1. Musculo-Skeletal Model

The musculo-skeletal model consists of two hind limbs attached to a trunk segment in the sagittal plane (Fig. 1-a). Each limb has thigh, shank, foot and toe segments connected by the hip, knee, ankle, and metatarsophalangeal (MTP) joints. The first three joints are actuated by six Hill-type muscles, which produce force based on their current length and velocity as well as their muscle stimulation, *s_m_* (muscle dynamics implemented as in Geyer and Herr, 2010). The muscles represent the illiopsoas (IP), gluteus maximus (GLU), vasti group (VAS), biceps femoris (BF), tibialis anterior (TA), and flexor digitorum longus (FDL) of an adult rat. In addition, the MTP joint is actuated by a passive spring and damper. All musculoskeletal parameters including segment dimensions, joint limits, muscle origins and insertions as well as muscle dynamics parameters are inspired by the anatomical studies of Charles et al. (2016) and Johnson et al. (2008, 2011) (see appendix A for details).

### 2.2. Treadmill and BWS System

The treadmill is modeled as an infinite surface that slides with a horizontal speed **v**_*b*_. The foot segments of the musculoskeletal model interact with this surface through frictional contact points (green circles in Fig. 1-a), which produce ground reaction forces, **F**_*gr*_, and are modeled as in Geyer and Herr (2010).

The BWS system mimics the system described in Nessler et al. (2005), whose long support lever and stiff harness effectively restrain the trunk from moving back and forth. The model accounts for this setup with a prismatic joint attached to the trunk that admits only vertical motion (Fig. 1-a). The joint is actuated by a constant suspension force, **F**_*bws*_, which can be adjusted for different levels of support.

### 2.3. Spinal Control Circuitry

Similar to our previous work (Sar and Geyer, 2020), we model spinal locomotion control as a large network of interconnected muscle reflexes. Going beyond this precedent, however, we include interlimb reflexes and separate controllers for stance and swing. For an example, the signal flow of one muscle reflex within this large network is illustrated in figure 1-b. A proprioceptive signal *p_ik_* originating from one muscle (*k*) connects in the spinal cord to two interneurons (IN_*st/sw*_) that belong to separate spinal controllers for stance and swing. The outputs of these two control INs converge onto the same alpha motoneuron (*α*MN), which innervates another muscle (*l*) with stimulation *s_l_*. Note that the two IN outputs are alternately suppressed depending on whether the stance or swing control of the corresponding leg is active. In total, the spinal control network has 1728 such muscle reflexes, based on 6 proprioceptive signals originating from each of the rat model’s 12 muscles and routing through all 24 control INs (two per muscle).

The implementation of the spinal control network uses simple elements. First, the proprioceptive signals are implemented as in Sar and Geyer (2020) and comprise the time-delayed (Δ_*k*_) muscle force 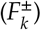, length 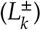, and contraction velocity 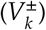, with the delay accounting for the time it takes neural signals to travel between muscles and the spinal cord, and the ± sign indicating excitatory or inhibitory projections. Second, the control INs are implemented as weighted sums,

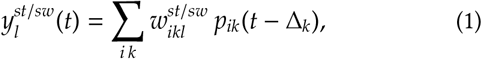

where the weights 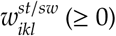 represent the synaptic strengths of the 72 projections that each IN receives (Fig. 1-c). Third, the αMNs are implemented as summation nodes whose outputs are saturated to ensure muscle stimulations between zero and 100%,

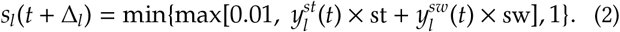

Finally, the switching between stance and swing control is achieved by contextual signals (Dietz, 2002; Yakovenko et al., 2004; Ekeberg and Pearson, 2005). In particular, swing phase is initiated when hip and ankle joint angles exceed threshold, which triggers activation of flexor muscles. Extensor muscles are later activated when ground foot contact sensory pathways are activated, which activates extensor muscles and results in body weight support. In addition, to ensure inter-limb coordination and body weight support, swing phase is only initiated when the other leg is touching the ground.

*Before* simulating spinal cord transection, we tune the muscle reflexes to produce ‘normal’ walking in the rat model. To this end, we hand-design muscle stimulation patterns that make the hind limbs walk on the treadmill at *ν_b_* = 5 cm s^-1^ without active support force, *F_bws_* = 0 N, and then tune the synaptic weights of the control INs by gradient descent until the muscle reflex control reproduces this locomotion behavior (see appendix B for details).

### 2.4. Spinalization and Gait Training Protocol

We model the effect of spinalization on the spinal control circuitry with an average reduction in the synaptic weights of all control INs. Specifically, the weights at the onset of the rehabilitation training are given by

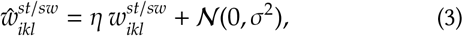

where *η* = 0.6 defines a 40% average reduction for all weights from before the spinalization, and the added Gaussian noise, 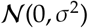, with the standard deviation *σ* set to 10% of the maximum weight, 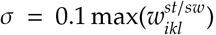, introduces some random variability about this average for each individual weight.

Although (3) oversimplifies the effects of spinalization, the scarce and, to an extent, uncertain knowledge about them precludes a finer interpretation. In general, what is known points to an overall reduction in the activity of the spinal cord circuitry (Côté et al., 2017). For instance, the activity of motoneurons below the site of the lesion is known to reduce dramatically, because neuromodulation is lost (Hounsgaard et al., 1988) and because excitatory projections onto these motoneurons decay in number (Witts et al., 2014) and strength (Petruska et al., 2007). It is further suggested that interneuron activity reduces as well (Kapitza et al., 2012), although some evidence points to the contrary due to diminished presynaptic inhibition from higher centers (Caron et al., 2020). Lastly, it is known that some detrimental effects of spinalization can be reversed in part by electrical stimulation and the use of neurotransmitters; specifically, in spinalized rats, combining both interventions can restore motoneuron output to about 60% of the pre-injury level (Courtine et al., 2009). The spinalization model (3) thus captures the net effect of an overall reduced spinal activity that is partially restored by intervention (*η* = 0.6), while at the same time acknowledging the uncertainty about changes in individual circuits 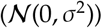.

Similar to the rat experiments described in Courtine et al. (2009), the damaged spinal control network in our model still produced rudimentary stepping on the treadmill when using a body weight support of *F_bws_* = 1N (about 50% BWS). We then followed this experimental study and simulated treadmill training at a belt speed of *ν_b_* = 7 cms^-1^. During training, we gradually reduced the BWS as the model’s muscle activities improved until full load-bearing by the animal model was reached (*F_bws_* = 0N, if applicable), and the synaptic weights of the spinal circuitry, 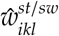, changed according to Hebbian learning models described in more detail below (section 2.5). We stopped the training once every weight changed less than 5% between consecutive strides.

### 2.5. Synaptic Adaptation during Gait Training

To address functional reorganization, we adapt the synaptic weights of the spinal circuitry using Hebbian learning strategies. Specifically, we build on the often applied Hebbian covariance learning (Sejnowski, 1977; Brown et al., 1990; Gerritsen et al., 1995; Abbott and Nelson, 2000; Bi and Poo, 2001; Brown and Milner, 2003; Legenstein et al., 2010; Markram et al., 2011). In this strategy, the weight of a synaptic connection between two neurons adapts as

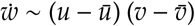

where *u* and *v* are the output signals of the pre- and postsynaptic neuron, respectively, and *ū* and 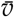 are their corresponding moving averages. Applying this learning strategy to our spinal control circuitry, we use

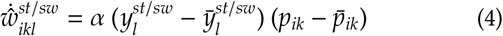

for the change of each synaptic weight 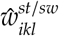 during gait training, where *α* is a constant learning rate (typically we use *α* = 0.1 – 0.5). A circuit interpretation of this weight adaptation is summarized in figure 2-a. Equation (4) can be interpreted as applying gradient *ascent* of the cost function

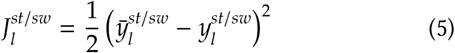

**Figure 2.**
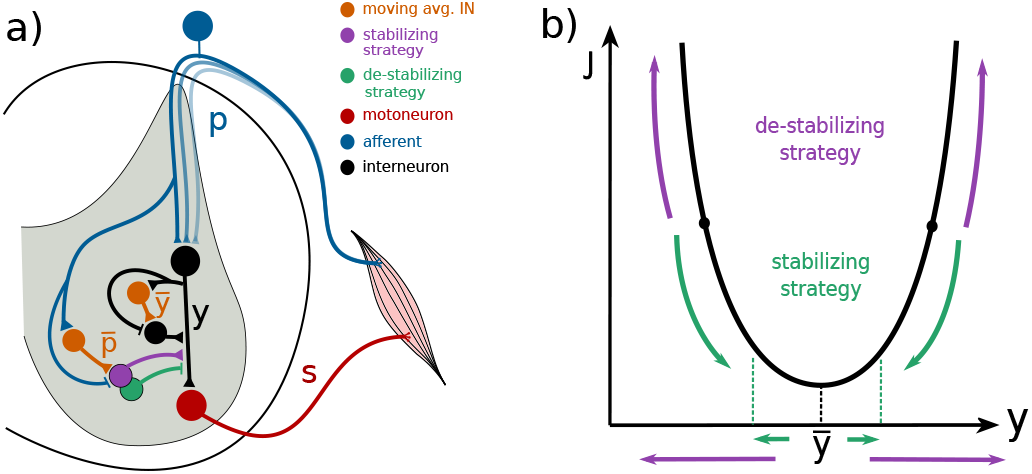
Synaptic weight adaptation during gait training. **a)** Potential circuits for implementing equations (4) and (6) of the synaptic weight adaptation at the spinal interneurons (compare Fig. 1). **b)** Cost function interpretation of the destabilizing and stabilizing weight adaptation strategies.

Consequently, the weight adaptation (4) is destabilizing; it will drive the spinal circuit activity away from its historical activity (Fig. 2-b) and, over time, into saturation (see appendix C for more details). By contrast, the alternative adaptation

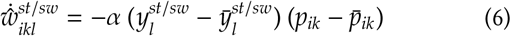

corresponds to gradient *aescent* of (5) and drives the activity of the spinal circuitry toward its recent average, converging eventually.

We will use the two learning strategies as simplified examples for a destabilizing (4) and a stabilizing (6) circuit adaptation during gait training, and examine the effects they have on the animal model’s functional recovery.

## 3. Results

As expected, the two alternative learning strategies lead to starkly different synaptic weight distributions when simulating gait training with the protocol described in section 2.4. Starting from the same initial conditions (Eq. 3), the synaptic weights of the two learning strategies express qualitatively and quantitatively different neural circuits at the end of the training. As an example, the resulting weight distribution at the stance control IN of the left leg TA is shown in figure 3. The destabilizing strategy leads to an almost binary distribution (Fig. 3-a), with the synaptic weights either growing unboundedly (mostly for the positive feedbacks, *F*^+^, *L*^+^, *L*^+^) or going to zero (mostly for the negative feedbacks, *F*^-^, *L*^-^, *L*^-^). By contrast, the weights of the stabilizing strategy converge to intermediate values for all pathways (Fig. 3-b).

**Figure 3.**
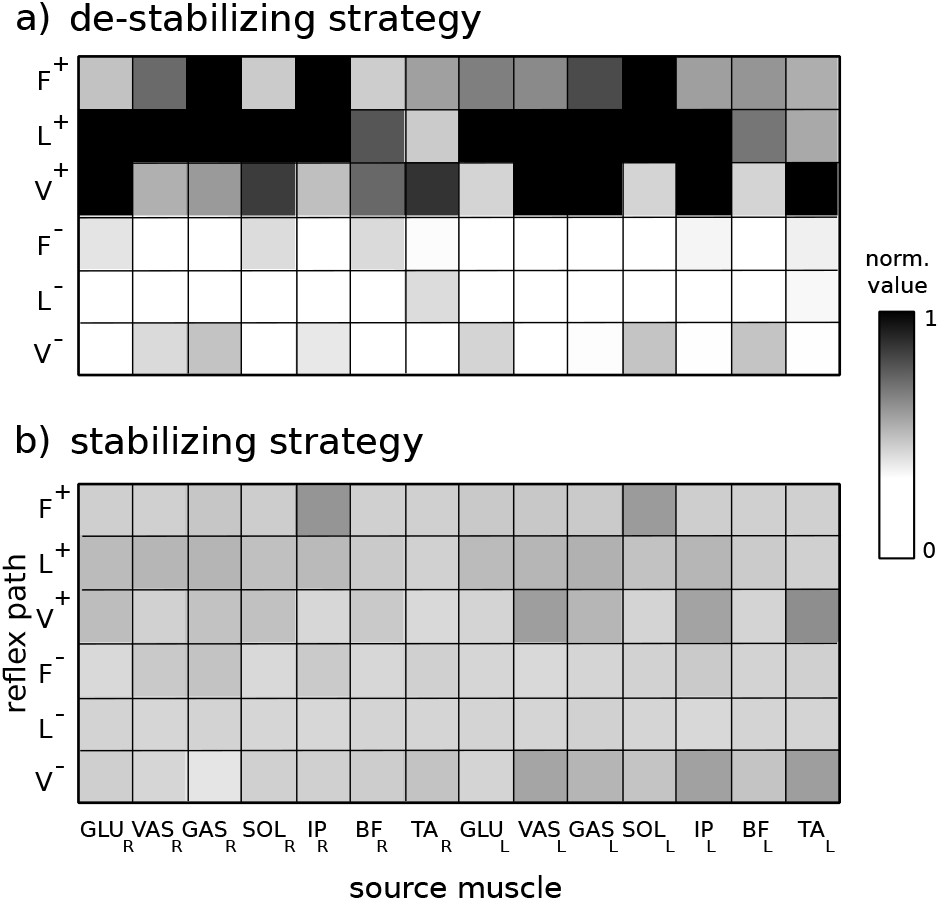
Example of resulting distribution of synaptic weights after gait training of spinal rat model with the destabilizing (a) and stabilizing (b) learning strategies. Shown are synaptic weights at stance control IN of the left leg TA muscle, normalized to the maximum weight of both panels.

Despite the stark difference in the resulting neural circuitry, both learning strategies lead to locomotion recovery as seen in experiments. For either strategy, gait training results in rhythmic locomotion of the spinal rat model inside the training apparatus with no additional BWS (*F_bws_* = 0 N). This result holds not only for walking at the specific training speed of the belt (Figs. 4) but extends to walking forward and backward at different speeds (see supplementary video), echoing the behaviors observed after gait training in animal experiments (Rossignol et al., 2001; Courtine et al., 2009).

**Figure 4.**
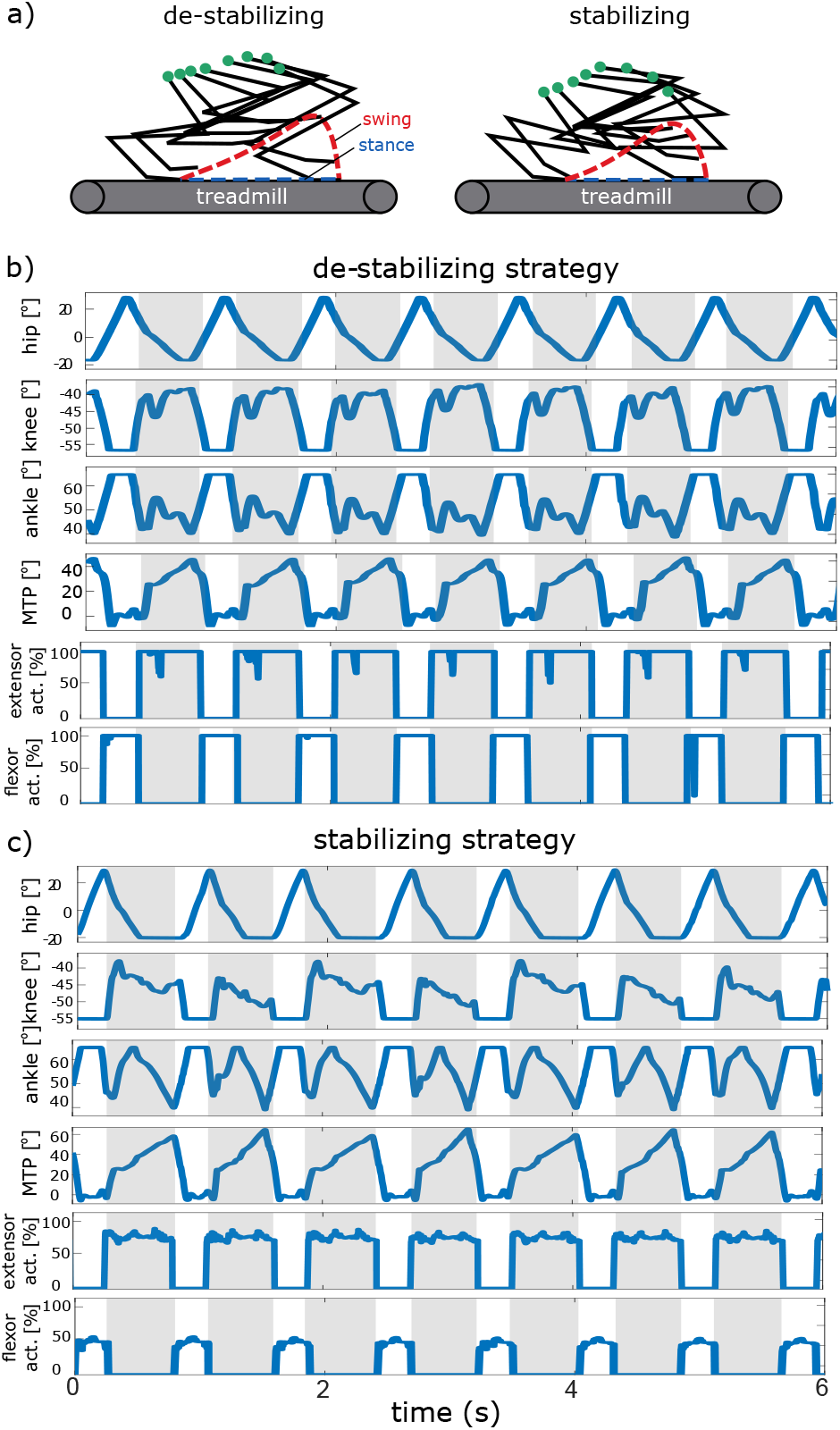
Example of model behavior after gait training. For both the destabilizing and stabilizing learning strategies, the spinal animal recovers walking (a) with rhythmic patterns of joint angles and muscle activations during stance (shaded areas) and swing (b, c). Notes: Treadmill belt speed set to 7cms^-1^.

When comparing the two learning strategies at the level of the specific outcome metrics reported in Courtine et al. (2009), we find that gait training with the destabilizing strategy more closely matches experimental observations (Fig. 5). Both learning strategies produce nearly identical changes for the gait timing, inter-limb coordination, weight bearing capacity, and limb endpoint trajectory (Fig. 5-a,b,c,d). However, substantial differences occur for the changes in muscle activities. While the stabilizing learning strategy returns the flexor and extensor amplitudes to their pre-injury value, the destabilizing strategy leads to excessive muscle amplitudes as observed in experiments (Fig. 5-e,f).

**Figure 5.**
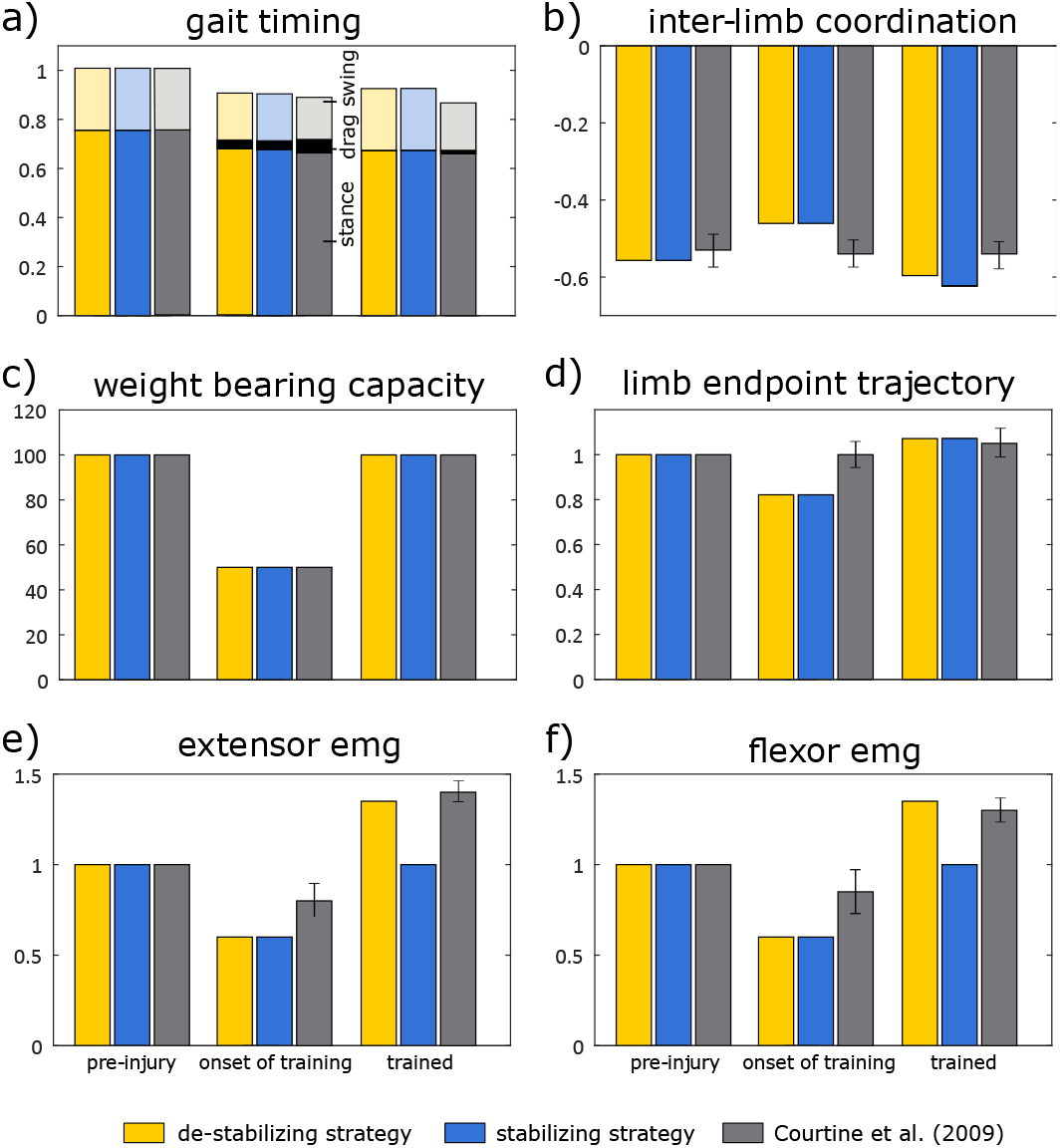
Model comparison at the level of outcome metrics reported in Courtine et al. (2009). (**a**) Gait timing: relative duration of stance, swing and foot drag. (**b**) Inter-limb coordination: cross-correlation between right and left leg movement. (**c**) Weight bearing capacity: percentage of body weight the animal model can support as it performs ten or more successful steps. (**d**) Limb endpoint trajectory: maximum step height of MTP joint above treadmill surface. (**e,f**) Extensor and flexor EMG: average muscle activity amplitudes. Notes: Experimental data replicated from ‘voluntary’ (pre-injury), ‘full combination’ (onset of training), and ‘trained with full combination’ (trained) conditions of figure 2 in Courtine et al. (2009). For simulation model, all metrics calculated as described in Courtine et al. (2009). Variability and kinematic similitude metrics reported in that paper omitted here, as description of their calculation appeared too vague.

## 4. Discussion

We used computer simulations to investigate if the functional reorganization observed in gait training of spinal animals leads to stable circuit function of locomotor control. Specifically, we developed a neuromuscular model of a spinalized rat exposed to gait training on a treadmill within a body-weight support system (Fig. 1). During training, we induced functional reorganization of the model’s spinal control circuitry using two alternative Hebbian learning strategies, one designed to simply drive the circuitry into saturation (destabilizing, Eq. 4) and the other designed to return the circuitry to some memory of its pre-injury function (stabilizing, Eq. 6). Despite this stark contrast in the learning strategies, we found that both lead to functional recovery of the simulated spinalized rat as seen in experiments (Barbeau and Rossignol, 1987; Leon et al., 1998; Rossignol et al., 2001; Ichiyama et al., 2005, 2008; Lavrov et al., 2008; Courtine et al., 2009; Rossignol and Frigon, 2011) (Fig. 4). Moreover, we found that the only significant difference in training outcomes between the two strategies concerns muscle activations, which showed the excessive amplitudes observed in experiments for the destabilizing strategy (Fig. 5).

### 4.1. Does functional recovery of gait imply recovery of circuit function?

Our results suggest that the functional recovery of gait observed in spinalized rat experiments does not necessarily imply recovery of circuit function, which should at least stabilize. If anything, the results for the model with the destabilizing learning strategy seem to agree more with the experimental observations, as not only the excessive muscle amplitudes more closely align but also the resulting large synaptic weights (Fig. 3-a) fit the increased H-reflex responses observed after training in spinalized rats (see, for instance, figure 5 in Courtine et al., 2009). Whether this conclusion truly applies to the recovery of spinalized rats remains open, however. Our simulation model clearly oversimplified the animal system, especially, its spinal circuitry and circuit reorganization during gait training. On the other hand, the simulations highlighted that the gait training protocols used in experiments might oversimplify what constitutes functional recovery of locomotion. The mechanical restrictions by the treadmill and body weight suspension system used in these protocols (Rossignol et al., 2001; Courtine et al., 2009; Rossignol and Frigon, 2011) largely determine the motion of the spinalized rat, and any enhancement of flexor and extensor activities during swing and stance, respectively, will likely generate locomotion and be perceived as recovery of gait.

### 4.2. Implications for Human Locomotor Training

While a large body of research is concerned with identifying the mechanisms that underlie functional reorganization (Kiehn, 2016; Smith and Knikou, 2016), the question of whether functional reorganization in the absence of supraspinal input produces stable circuit function is not commonly explored. Answering it will have implications for human gait rehabilitation. It is well established that gait training triggers spinal circuitry reorganization not only in incomplete spinal cord injured patients but also in complete ones (Smith and Knikou, 2016). But there are clear differences between the two groups. For instance, in complete spinal cord injured patients the soleus H-reflexes cannot be conditioned (Knikou and Mummidisetty, 2014), voluntary muscle control is severely impaired, and gait phase-dependent EMG activity remains absent even after training (Smith and Knikou, 2016). The differences may motivate the exploration of group-specific rehabilitation protocols (Eisdorfer et al., 2020). However, if functional reorganization in the absence of supra-spinal input is bound to drive the circuitry into saturation, then gait training is unlikely to recover locomotor function beyond overall muscle activity, regardless of rehabilitation protocol.

**Table 1.** Leg segment masses and lengths.

| segment | mass (g) | length (cm) |
| --- | --- | --- |
| trunk | 150 | 9.0 |
| thigh | 7 | 3.6 |
| shank | 5 | 4.1 |
| foot | 3 | 2.5 |

**Table 2.** Joint limits.

| joint | $\theta_{min}$ (deg) | $\theta_{max}$ (deg) |
| --- | --- | --- |
| hip | -20 | 50 |
| knee | -55 | 60 |
| ankle | -50 | 65 |
| MTP | -70 | 100 |

### 4.3. Telling apart stable and unstable recovery in experiments

One way of telling apart a stable from an unstable reorganization of the spinal circuitry could be to train spinalized rats at below full weight bearing capacity for much longer than usual. Because full weight bearing capacity is an essential goal in gait rehabilitation, this is not done typically. Most training protocols of spinal animals gradually reduce suspension until the animal recovers full body weight support, which is achieved after several weeks of training along with the recovery of gait (Rossignol et al., 2001; Rossignol and Frigon, 2011; Loy and Bareyre, 2019). If full support strains what the animal muscles can provide, it will not be possible to tell a difference between the stable and unstable reorganization of the spinal circuitry. However, when training at support levels well below muscle capacity, the two types of reorganization should manifest in different animal behavior over time. A stable reorganization should lead to animal walking as known from gait training, whereas an unstable reorganization should manifest as walking with an over-powered, jumping-like gait due to eventual circuit saturation.

## Appendix

### A. Musculo-skeletal Model Parameters

Tables 1 to 4 summarize the musculo-skeletal parameters of the rat model. They are based on anatomical studies (Johnson et al., 2008, 2011; Charles et al., 2016) with slight modifications to account for model simplifications.

### B. Tuning of Reflex Circuitry before Spinal Transection

We used stochastic gradient descent to tune the muscle reflex network and produce ‘normal’ walking in the rat model. First, we hand-designed reference stimulation patterns 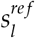 for the model’s extensor (e) and flexor (f) muscles based on sinusoids,

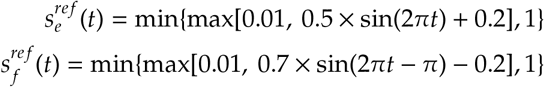

tuned to ensure the hind limbs walk on the treadmill at *v_b_* = 7 cm s^-1^ without active support force, *F_bws_* = 0 N. Similar to Sar and Geyer (2020), we then used stochastic gradient descent of the cost functions 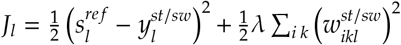 to learn the muscle reflex control that reproduces this walking behavior. More specifically, we initialized all the synaptic weights 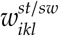 of the control INs (Eq. 1) to zero and then updated them with the rule

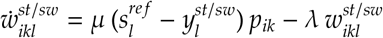

during repeated treadmill walking (learning and regularization rates set to *μ* = 0.1 and λ = 0.01, respectively). Finally, we stopped this process once the weights converged and confirmed the resulting muscle-reflex network produces treadmill walking in the rat model.

**Table 3.** Muscle moment arms.

| muscle | $d_{hip}$ (mm) | $d_{knee}$ (mm) | $d_{ankle}$ (mm) |
| --- | --- | --- | --- |
| IP | 6.0 | - | - |
| GLU | -6.0 | - | - |
| BF | -1.0 | -6.0 | - |
| VAS | - | 3.5 | - |
| GAS | - | -1.0 | -1.0 |
| TA | - | - | 6.0 |
| SOL | - | - | -4.0 |

**Table 4.** Hill-type muscle parameters.

| muscle | $F_{max}$ (N) | $l_{opt}$ (mm) | $l_{slack}$ (mm) | $v_{max}$ (mm s<sup>-1</sup>) |
| --- | --- | --- | --- | --- |
| IP | 60 | 20.9 | 3.9 | 1200 |
| GLU | 60 | 27.4 | 11.2 | 1200 |
| BF | 45 | 14.3 | 18.6 | 900 |
| VAS | 60 | 24.5 | 8.7 | 900 |
| GAS | 60 | 15.3 | 29.9 | 1200 |
| TA | 60 | 9.3 | 19.2 | 1200 |
| SOL | 50 | 9.2 | 21.3 | 1200 |

### C. Cost Function Interpretation of Learning Strategies

With the average interneuron output defined as

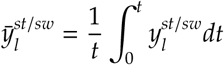

the gradient of the cost function (5) with respect to the weight 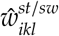 resolves to

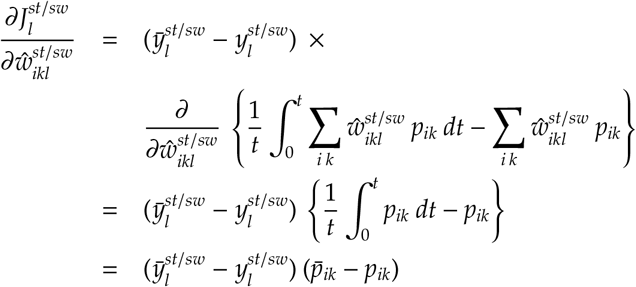

Hence, a weight change 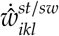 defined as in (4) and (6) corresponds to gradient ascent and descent of the cost function (5), respectively (Fig. 2-b). In the first case, the weight change will drive the IN output 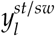 away from its recent average, 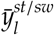, causing individual weights 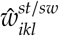 to approach ever larger values or zero over time. In the second case, the IN output and its recent average will converge to each other, stabilizing the circuitry.

